# An atlas of silencer elements for the human and mouse genomes

**DOI:** 10.1101/252304

**Authors:** Naresh Doni Jayavelu, Ajay Jajodia, Arpit Mishra, R. David Hawkins

**Affiliations:** Division of Medical Genetics, Department of Medicine, Department of Genome Sciences, Institute for Stem Cell and Regenerative Medicine, University of Washington School of Medicine, Seattle, WA USA

## Abstract

The study of gene regulation is dominated by a focus on the control of gene activation or controlling an increase in the level of expression. Just as critical is the process of gene repression or silencing. Chromatin signatures have allowed for the global mapping of enhancer *cis*-regulatory elements, however, the identification of silencer elements by computational or experimental approaches in a genome-wide manner are lacking. We present a simple but powerful computational approach to identify putative silencers genome-wide. We used a series of consortia data to predict silencers in over 100 human and mouse cell or tissue types. We performed several analyses to determine if these elements exhibited characteristics expected of a silencers. Motif enrichment analyses on putative silencers determined that motifs belonging to known transcriptional repressors are enriched, as well as overlapping known transcription repressor binding sites. Leveraging promoter capture HiC data from several human and mouse cell types, we found that over 50% of putative silencer elements are interacting with gene promoters having very low to no expression. Next, to validate our silencer predictions, we quantified silencer activity using massively parallel reporter assays (MPRAs) on 7500 selected elements in K562 cells. We trained a support vector machine model classifier on MPRA data and used it to refine potential silencers in other cell types. We also show that similar to enhancer elements, silencer elements are enriched in disease-associated variants. Our results suggest a general strategy for genome-wide identification and characterization of silencer elements.

## INTRODUCTION

It has long stood that repressive transcription factors can bind to the promoters of genes through silencer elements to inactivate gene expression ^1–4^. In 1985, the yeast mating type loci revealed that distal silencer elements could control gene expression from afar ^5^. The role for distal silencer elements in mammals was demonstrated shortly thereafter through a silencer element located several kilobases upstream of the rat insulin gene ^6^. It would be a decade a later before key experiments identifying a silencer in the intron of the mouse and human *CD4* genes would revealed the important role that silencers can play in lineage specificity and cell fate determination, as this silencer represses *CD4* expression in CD8+ T cells ^7,8^. Later several studies identified genomic sequences with silencer properties that are opposite of enhancers across many species ^9–13^. While dozens of mammalian silencers have been identified, these elements are largely understudied, possibly due to our biased focus on gene upregulation and the clandestine nature of those with non-promoter locations.

Outside of promoter regions, silencers along with enhancers and insulators create a complex array of distal *cis*-regulatory elements. These elements, and the factors that bind to them, are important for the nuanced output of RNA levels across cell types. Identification of distal *cis*-regulatory elements and our understanding of the regulation of gene expression in mammalian genomes has been greatly facilitated by the genome-wide mapping of element-specific histone modifications or transcription factors, e.g., histone H3 lysine 4 monomethylation (H3K4me1) distinguishing enhancers ^14^ and CTCF binding to a subset of putative insulators ^15^. While the presence of repressor sequences within *cis*-regulatory elements such as promoters has recently become more evident by tiling millions of oligos across these regions and testing by massively parallel reporter assays (MPRA) ^16^, distinct annotations for distal silencer elements in the human and mouse genomes are still missing from our *cis*-regulatory lexicon.

Our goal was to identify distinct silencer elements distal to the genes they regulate. Implementing a simple subtractive analysis approach based on DNase hypersensitive sites (DHS) and known other regulatory elements results in an atlas of more than 1.5 million unique putative silencer elements in the human genome and a second atlas of ∼1 million in the mouse genome. We find that silencers are enriched for motifs of known repressive transcription factors, and *de novo* motifs for potential cognate transcriptional repressors. We validated our predictions by integration of ChIP-seq data for known repressors, direct silencer interactions with repressed target genes, and testing via MPRA. Based on our MPRA data for 7,500 putative silencers, we trained a support vector machine (SVM) classifier to further refine silencer element predictions in the human (and mouse) genomes. Putative silencers are often enriched for disease associated variants in expected cell types or lineages. These atlases, which will require more intensive study and validation from the field, should aid our understanding of gene expression through negative regulation of expression.

## RESULTS

### Simple subtractive analysis approach predicts genome wide putative silencer elements

We devised an efficient simple subtractive analysis (SSA) approach to predict genome-wide putative silencer elements (Fig. 1a). The potential *cis*-regulatory elements should present in open chromatin. We can generate open chromatin data either from DNase-seq (DNase I hypersensitive sites sequencing) or from ATAC-seq (Assay for Transposase-Accessible Chromatin using sequencing) and other *cis*-regulatory elements data from ChIP-seq (chromatin immunoprecipitation coupled with sequencing) for any cell type or organism. In this SSA approach, we subtract enhancers (H3K4me1 peaks), promoters (H3K4me3 peaks), potential insulators (CTCF peaks) and active TSS from open chromatin (DHS) and assign the leftover (or remaining) DHS as silencers, we term these elements as “putative silencer elements”.

**Figure 1.**
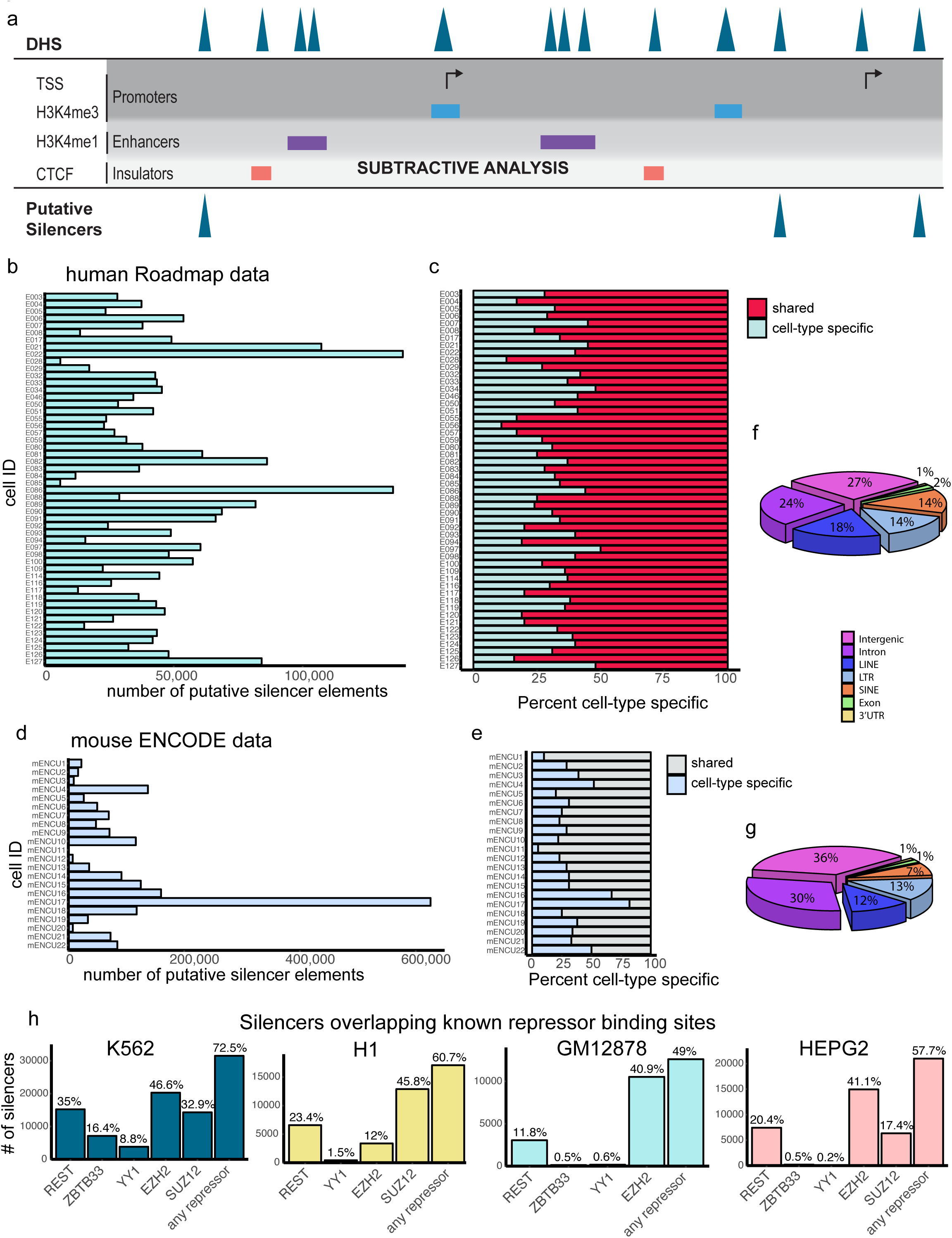
Overview of predicted putative silencer elements in human and mouse. **(a)**Schematic of the simple subtractive analysis (SSA) approach. Bar plots presenting the count of predicted silencer elements across different cell types and tissues in human from Roadmap (b) and mouse genomes from ENCODE (d). Distribution of cell-type specific silencer elements in human (c) and mouse genomes (e). Pie chart showing the genomic distributions of putative silencer elements in human (f) and mouse (g). (h) Bar plots presenting the count of silencer elements at known repressor TFBS based on ChIP-seq data in K562, H1, GM12878 and HEPG2 cell types.

Using this SSA approach we predicted 2,315,105 putative silencer elements in human genome spanning across 82 cell types from the Roadmap ^17^ and ENCODE consortia ^18^, and 1,299,866 elements in the mouse genome for 22 cell types from the ENCODE consortium 18 (Fig. 1b,d and Supplementary Fig. 1a,b and Supplementary Table 1). On average ∼52,360 elements and ∼87,800 elements percell type were identified in human and mouse genomes, respectively. We further filtered to identify cell-type specific (unique) putative silencer elements. In total 1,534,618 unique putative silencers in human cell types and 984,477 unique ENCODE consortium ^18^ (Fig. 1b,d and Supplementary Fig. 1a,b and **Supplementary Table 1**). On average ∼52,360 elements and ∼87,800 elements per elements are predicted in mouse cell types (Fig. 1c,e and Supplementary Fig. 1c). The identified putative silencer elements are largely present at intergenic, introns and repeat elements of the genome for both human and mouse (Fig. 1f,g) and located relatively close to gene TSS (Supplementary Fig. 1d,e).

### Putative silencer elements are enriched with known repressor TFBS

Transcription factors (TFs) bind at *cis*-regulatory elements and regulate gene expression in response to external cues. Activator TFs bind at enhancers to enhance expression and repressor TFs bind at silencer elements and repress gene expression. Based on this notion, we determined whether SSA-predicted silencers are enriched with repressor TFBS. We intersected putative silencer elements with known repressor TFBS within the same cell type using ChIP-seq data for REST, YY1, ZBTB33, SUZ12 and EZH2. We found that 73%, 61%, 49% and 58% of total putative silencers in K562, H1, GM12878, HEPG2 cells, respectively, are enriched with one of the known repressor TFBS (Fig. 1h). This corroborates a likely role for the putative silencers in negative regulation of gene expression.

### Characterization of putative silencer elements

Given the limited availability of ChIP-seq data for transcriptional repressors, we performed motif enrichment analysis on all cell types to identify other potential binding factors. As expected motifs belonging to REST are highly enriched in putative silencers of all 52 tested human cell types from Roadmap (Fig. 2a,b). Surprisingly, motifs belonging to repressors ZBTB33 and YY1 are only enriched in 9 and 3 cell types, respectively. Motifs of USF1, BATF, BACH2, FRA1, ATF3, FOSL2, JUN, NRF2, NFE2 and RFX family of TFs are also enriched at silencers of many cell types. Previous studies reported that many of these TFs display repressor activity 19–22.

**Figure 2.**
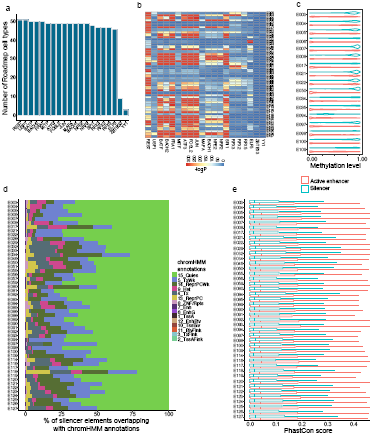
Characteristics of putative silencer elements. **(a)** Bar plot presenting the count of cell types enriched with TF motifs across 52 cell types and tissues from Roadmap. (b) Heatmap of enrichment (-log(P-value)) of TF motifs across 52 cell types. (c) Violin plot showing the distribution of methylation levels at putative silencer elements and active enhancers across 17 cell types. (d) Bar plot showing the distribution of chromHMM annotations of putative silencer elements across 52 human cell types. (e) Box plot of PhastCon conservation scores for putative silencers and active enhancers across 52 cell types.

Next, we explored additional characteristics of the putative silencer elements. DNA cytosine methylation is an important epigenetic modification affecting gene expression patterns during development and disease. DNA hypermethylation at promoters and CpG Islands often results in repression of gene expression and hypomethylation results in activation. Similarly, enhancer elements are either largely unmethylated or lowly methylated ^23,24^. However, a number of transcriptional repressors are known to bind methylated DNA ^25–27^, suggesting some silencer elements would be methylated. We computed the average methylation levels for putative silencer elements using whole genome bisulfite sequencing (WGBS) data from 17 cell types and found them to largely be in a hypermethylated state (Fig. 2c and Supplementary Fig. 2a). We also compared the average methylation of active enhancers (DHS marked with H3K4me1 + H3K27ac) for corresponding cell types and found significantly (P-value < 2.2e-16) lower methylation than at putative silencers. Putative silencers are on average 75% methylated compared to 43% methylation in active enhancers across the 17 cell types (Supplementary Fig. 2a).

Enhancers and promoters can be annotated in the genome based on distinct chromatin signatures ^28^. Such a chromatin signature for silencers would facilitate their genomic localization. To investigate such, we made use of already available chromHMM annotations for 52 human cell types from the Roadmap consortium. Unfortunately, the majority of the putative silencer elements belong to a largely uncharacterized annotation category or lack of known histone mark signal (Quies) (Fig. 2d and Supplementary Fig. 2b). This uncharacterized category varies from 21% to 80% across cell types. The next three categories are weak transcription (TxWk) and weak Polycomb Repressor Complex (ReprPCWk) followed by heterochromatin (Het). While limited, all three categories are consistent features of transcriptional repression and some repressor TFs.

Lastly, we examined conservation scores to check whether putative silencer elements are conserved in vertebrate evolution. We used the 100-way phastCons score comparing the human genome to 99 other vertebrate genomes. The computed phastCons scores suggest that silencer elements are less or non-conserved across species similar to enhancer elements ^29^ (Fig. 2e). The average phastCon score for putative silencers is 0.12 where as for active enhancers it is 0.14 across 52 cell types.

### Putative silencers interact with inactive and lowly expressed genes

Previous promoter capture HiC (p-CHiC) studies reported that transcriptionally inactive or lowly expressed genes promoters are interacting with uncharacterized regions of genome, which suggest that they may act as silencers ^30,31^. We investigated whether our predicted putative silencer elements are interacting with inactive genes. For this analysis we obtained four p-CHiC data sets, two for each in human (GM12878 and CD34+ cells) and mouse (mESCs and mouse fetal liver cells (FLCs)) cell types ^30,31^. We overlapped putative silencer elements with non-baited fragments (or promoter interacting fragments) to find their interacting genes. In total, we find 19,333 putative silencers are interacting with 19,362 genes in GM12878 cells and 11,093 putative silencers are interacting with 11,654 genes in CD34+ cells. We find 5,305 and 2,198 inactive genes (with RPKM = 0) are interacting with 13,159 and 3,719 putative silencers in GM12878 and CD34+ cells, respectively (Fig. 3a,d and Supplementary Fig 3a,b). An additional set of 6,624 and 4,044 lowly expressed genes (with RPKM between >0 and 2) are interacting with 15,038 and 6,879 putative silencer elements in GM12878 and CD34+ cells (Fig. 3a,d and Supplementary Fig 3a,b). Furthermore, the overall expression of all genes interacting with putative silencers showed significantly lower expression than genes interacting with active enhancers in both GM12878 and CD34+ cells (Fig. 3b,e). Next, we looked at the chromatin states of transcription start sites (TSS) of genes interacting with putative silencers categorized into different expression ranges (Fig. 3c,f). The majority of inactive and lowly expressed gene TSS are uncharacterized with no known chromatin mark (Quies: GM‐ 50%; CD34‐ 53%) and to some extent enriched with Supplementary Fig 3a,b). An additional set of 6,624 and 4,044 lowly expressed weak polycomb repressor complex (ReprPCWk: GM‐ 21%; CD34‐ 16%), weak transcription (TxWk: GM‐ 9%; CD34‐ 6%), followed by heterochromatin (Het: GM‐ 3%; CD34‐ 3%). These chromatin states might be expected for genes repressed through distal silencers. In contrast, highly expressed genes are enriched with active TSS (TssA: GM‐ 43%; CD34‐ 30%) and transcription (Tx: GM‐ 20%; CD34‐ 30%) annotation categories. The fraction of uncharacterized and active TSS are in opposing trends with gene expression (Fig. 3c,f).

**Figure 3.**
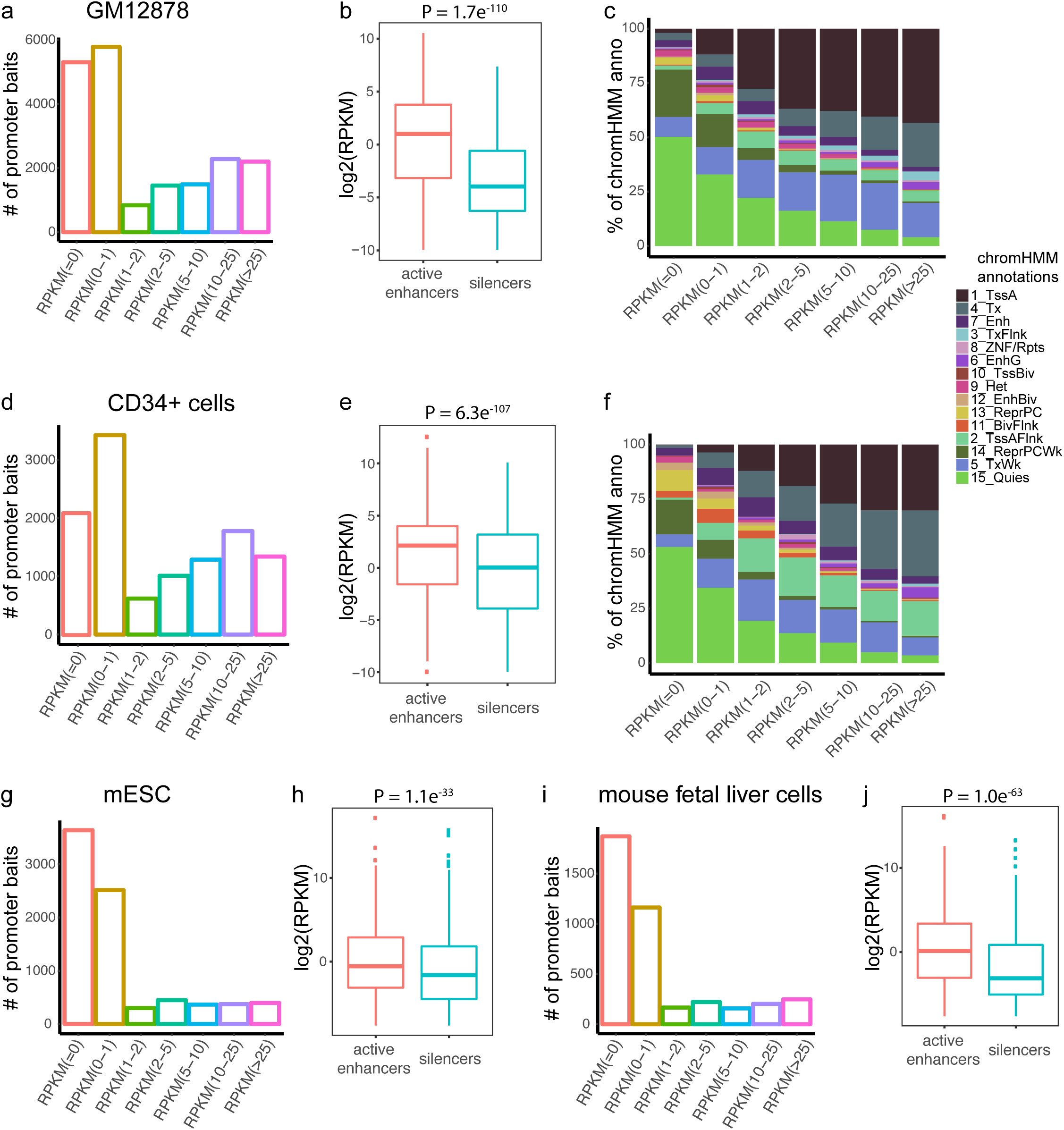
3D Genome interactions of putative silencer elements. **(a)** Bar plot presenting the counts of genes expressed at different expression levels (RPKM) interacting with putative silencer elements in GM12878 cells. (b) Box plot comparing the expression levels (RPKM) of genes interacting with putative silencer elements and active enhancers in GM12878 cells. (c) Bar plot showing the distribution of chromHMM annotations for TSS of genes interacting with putative silencer elements in GM12878 cells. (d) Bar plot presenting the counts of genes expressed at different expression levels (RPKM) interacting with putative silencer elements in CD34+ cells. (e) Box plot comparing the expression levels (RPKM) of genes interacting with putative silencer elements and active enhancers in CD34+ cells. (f) Bar plot showing the distribution of chromHMM annotations for TSS of genes interacting with putative silencer elements in CD34+ cells. (g) Bar plot presenting the counts of genes expressed at different expression levels (RPKM) interacting with putative silencer elements in mouse embryonic cells (mESCs). (h) Box plot comparing the expression levels (RPKM) of genes interacting with putative silencer elements and active enhancers in mESCs. (i) Bar plot presenting the counts of genes expressed at different expression levels (RPKM) interacting with putative silencer elements in mouse fetal liver cells (FLCs). (j) Box plot comparing the expression levels (RPKM) of genes interacting with putative silencer elements and active enhancers in FLCs.

We repeated the p-CHiC analysis with the mouse data sets and found more profound results (Fig. 3g-j). Briefly, a total of 13,560 putative silencers are interacting with 8,032 genes in mESCs and 3,656 putative silencers are interacting with 4,025 genes in FLCs. We find 6,155 and 3,032 genes with RPKM in the range of 0-1 are interacting with 11,459 and 3,050 putative silencers in mESCs and FLCs, respectively (Fig. 3g,i and Supplementary Fig 3c,d). Overall, 80% and 79% of total silencer interacting genes are within the expression range of 0-2 in mESCs and FLCs, respectively (Fig. 3g,i and Supplementary Fig 3c,d). Silencer interacting genes showed overall lower expression than enhancer interacting genes in both cell types (Fig. 3h,j). Altogether, this analysis provides strong evidence for silencer activity for the SSA-predicted putative silencer elements.

### Functional validation of putative silencer elements in K562 cells

Having shown that some putative silencer elements are demonstrating silencer activity through the direct interaction with repressed genes, we proceeded to functionally validate a subset of putative silencer elements and quantitate their activity via massively parallel reporter assays (MPRA) using the STARR-seq ^32^ approach in K562 cells (Fig. 4a). The human STARR-seq vector utilizes the Super Core Promoter (SCP1), which was designed and shown to be stronger than the CMV promoter ^33^, therefore, the GFP reporter expression should be susceptible to detectable decreases in expression. We selected 7,430 putative silencer elements, 20 known silencer elements ^16,34^, 20 known enhancer elements ^16^ and 20 randomly selected regions, which act as a control set (**Supplementary Table 2**). Oligos were designed to span 200nt of the silencer sequences flanked by 15nt adapter sequences and synthesized en masse. Of 7,430 selected putative silencer elements, 3,705 silencers have at least one TFBS belonging to well-known repressor TFs (REST, YY1, ZBTB33, EZH2 and SUZ12) and 3,725 silencers have at least one known motif belonging to GATA1, GATA2, GATA3, GATA4, BACH1, BACH2, TCF12, SMAD3, FLI1, RUNX1, KLF4, ZFP187, ZNF263, ZBTB7B and GFI1B (actual numbers are provided in **Supplementary Table 2**). All of which were reported to have repressor activity, while primarily serving as activators ^19,35–41^. As expected, enhancers showed the highest activity and known silencers the lowest activity, while putative silencers and control random regions showed similar level of activity (Fig.4b). We found 3,817 of the putative silencers (2,230 silencers with known repressor TFs status and 1,587 silencers with other TFs status) to have activity less than median activity level of control random regions at 10% FDR (Fig. 4c). Both categories of putative silencers, those with known repressor status and silencers with other TFs status, showed significantly lower activity than control random regions (Fig. 4d). We also looked at the distribution of TF status in order to identify strong candidate TFBS for potential silencer activity (**Supplementary Table 2**). Though the majority of TFBS showed 50% success rate, TFBS of EZH2, SUZ12, KLF4, RUNX1, TCF12, REST and YY1 would be strong candidates indicative of potential silencers. Because most known TFBS motifs are obtained from transcriptional activators, we also performed a *de novo* motif analysis. The *de novo* analysis identified several novel TFs motifs. When considering the closest related known motif, these belong to REST, YY1 and ZBTB33 (Fig. 4f).

**Figure 4.**
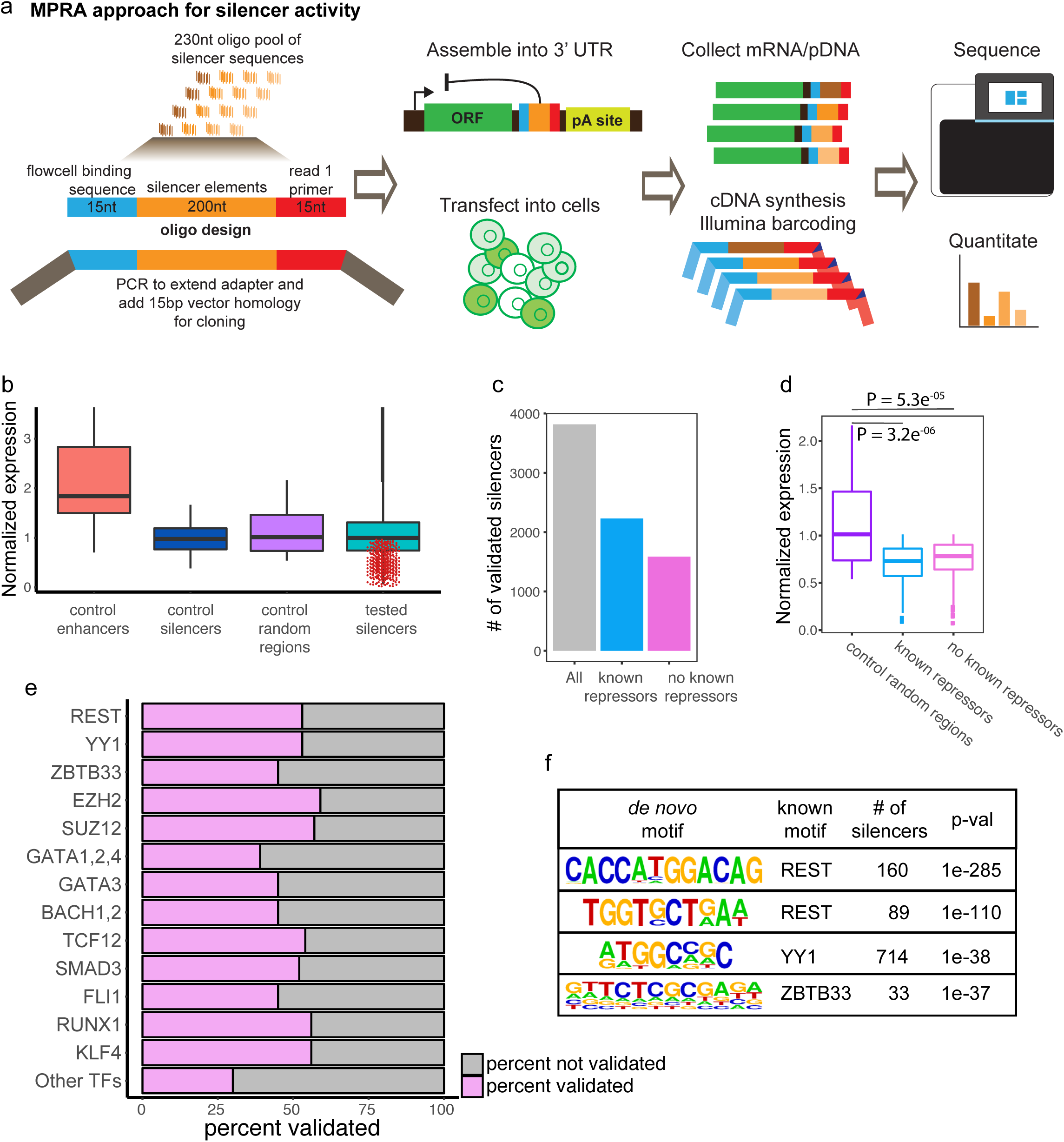
Functional validation of activity of putative silencer elements in K562 cells. **(a)** Schematic of MPRA/STARR-seq approach for measuring the activity of putative silencer elements. (b) Box plot displaying the activity level distributions of enhancers, control silencers, control random regions and tested putative silencer elements. Red dots denote the putative silencer elements showing silencer activity. (c) Box plot showing the count of all validated silencer elements, and counts of validated silencers overlapping with known repressor TFBS (TFBS belonging to REST, YY1, ZBTB33, SUZ12 and EZH2) or with other TFs motifs. (d) Box plot comparing the activity level distributions of control random regions and validated silencers categorized to overlap contain known repressor TFBS and other TFs motifs. (e) Bar plot showing the fraction of validated silencers categorized by different TFs. (f) Enriched *de novo* motifs related to well-known repressor TFs in validated silencers.

### SVM model development and predictions

Recent studies showed that well-trained support vector machine (SVM) models can predict cell-type specific *cis*-regulatory elements from a given set of nucleotide ^43^. Using a gapped k-mer SVM (gkmSVM) ^44,45^, we trained the classifier based on MPRA functional validation data to further refine our silencer predictions (Fig. 5a). We chose the top 2,200 putative silencer sequences with lowest activity as a positive set and the bottom 2,200 silencers with highest activity as a negative set for the gkmSVM model. We trained the gkmSVM model on 80% of the data and used the remaining 20% of data for testing the model. We checked the performance sequences ^42,^ of the model on test data by generating the receiver operating characteristic (ROC) curve by plotting true positive rate versus false positive rate and the precision recall curve (PRC). The model accurately predicted the positive silencers on the test set and the model performance in terms of area under the curve (AUC) for the ROC curve is 0.795 and for PRC is 0.797 (Fig. 5b,c). We then used the model for predicting potential silencers across all cell types from the list of human SSA-predicted putative silencers. We chose the threshold for the gkmSVM score where the model’s accuracy is maximum in order to classify the positive silencers from negative ones. The percent of gkmSVM positive predictions range from 28% to 74% for Roadmap human cell types and it varies from 34% to 78% for ENCODE human cell types (Fig. 5d,e and **Supplementary Table 3**).

**Figure 5.**
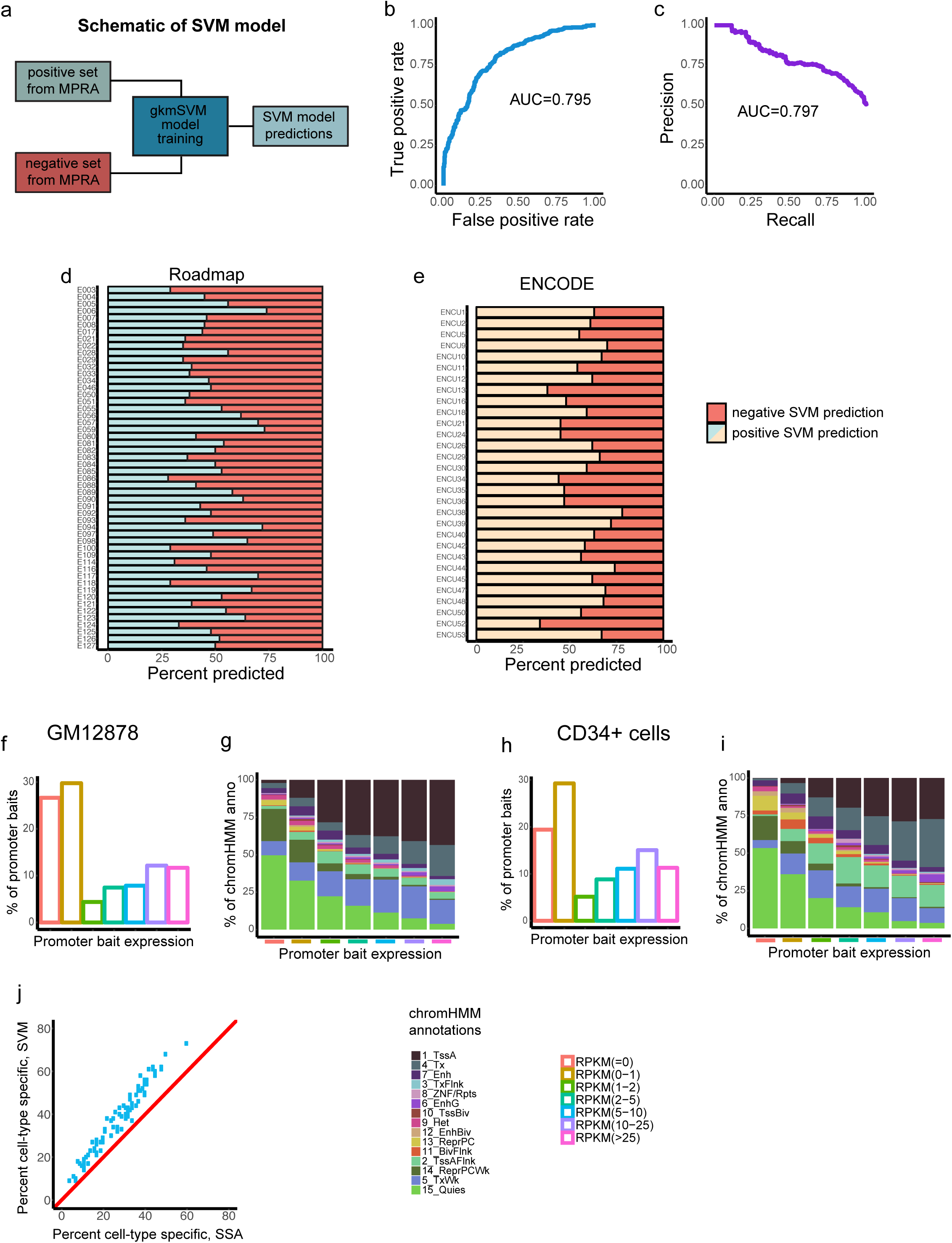
Overview of SVM model and silencer element predictions. **(a)** Schematic overview of the gkmSVM model for silencer predictions. The trained gkmSVM model performance on test data shown by receiver operating characteristic (ROC) curve (b) and precision recall curve (c). Bar plot showing the fraction of SVM-predicted silencers tested on SSA-predicted silencers in 52 human cell types and tissues from Roadmap (d) and 54 human cell types and tissues from ENCODE (e). (f) Bar plot presenting the counts of genes expressed at different expression levels (RPKM) interacting with SVM-predicted silencer elements in GM12878 cells. (g) Bar plot showing the distribution of chromHMM annotations for TSS of genes interacting with SVM-predicted silencer elements in GM12878 cells. (h) Bar plot presenting the counts of genes expressed at different expression levels (RPKM) interacting with SVM-predicted silencer elements in CD34+ cells. (i) Bar plot showing the distribution of chromHMM annotations for TSS of genes interacting with SVM-predicted silencer elements in CD34+ cells. (j) Comparing percent of cell type specific silencer elements identified from the SVM classifier and SSA approach. Each dot represent 82 human cell types and diagonal line represent reference for equal performance.

Next, we examined whether the SVM-predicted silencers can improve the fraction of their interacting genes with expression ranging from 0-2 RPKM compared to SSA-predicted silencers. We repeated the same analysis as in the previous section. Surprisingly, the SVM-predicted silencers have relatively similar fractions of interacting genes with RPKM of 0-2 to total interacting genes as did SSA-predicted silencers in both GM12878 (61% SVM vs 62% SSA) and CD34+ cells (54% SVM vs 54% SSA) (Fig. 5f,h and Supplementary Fig 4c,d). The chromatin status of TSS for interacting genes are also similar to what was observed for SSA-predicted silencers (Fig. 5g,i). Overall, the SVM classifier predicted relatively more cell-type specific silencers than SSA approach (Fig. 5j and Supplementary Fig 4a,b). This may aid future detection of cell-specific silencer elements when the entire repertoire of annotated elements needed to predict silencers is not available in a given cell or tissue type.

### Disease-associated variants are enriched at putative silencer elements

Several earlier studies showed that disease-associated single nucleotide polymorphisms (SNPs) from genome-wide association studies (GWAS) are prevalent at non-coding regions of the genome, specially at *cis*-regulatory elements, and often in enhancers of cells thought to be associated with the disease ^17,46–48^. We next investigated whether putative silencer elements were also enriched for disease-associated variants. First, we determined how many SNPs are present at silencer elements. For this, we downloaded all disease SNPs from the NHGRI-EGI GWAS catalogue. We included both lead SNPs and SNPs in linkage disequilibrium (LD) (r2 >= 0.8) for overlap with silencer elements. We found that 19,088 SNPs belonging to 1,500 disease traits are present at silencer elements across all cell types. Next, we examined significant enrichment of specific disease SNPs at silencers for each cell type. This analysis resulted in ∼25% of these diseases (369 out of 1,500) being significantly enriched (adjusted P-value < 0.01, hypergeometric test) at silencers for one or more cell types (Fig. 6a,b). Next, we asked whether these disease SNPs are enriched in relevant disease cell types. As a test case, we examined whether enrichment of autoimmune diseases are specific to blood cell types (Fig. 6c). Indeed, we found multiple sclerosis, rheumatoid arthritis, coronary artery disease, adult asthma, autoimmune hepatitis type 1, and coagulation factor levels to be specifically enriched at silencer elements of blood cell types. Overall, our results illustrate that putative silencer elements are also enriched with disease-associated SNPs similar to other *cis*-regulatory elements.

**Figure 6.**
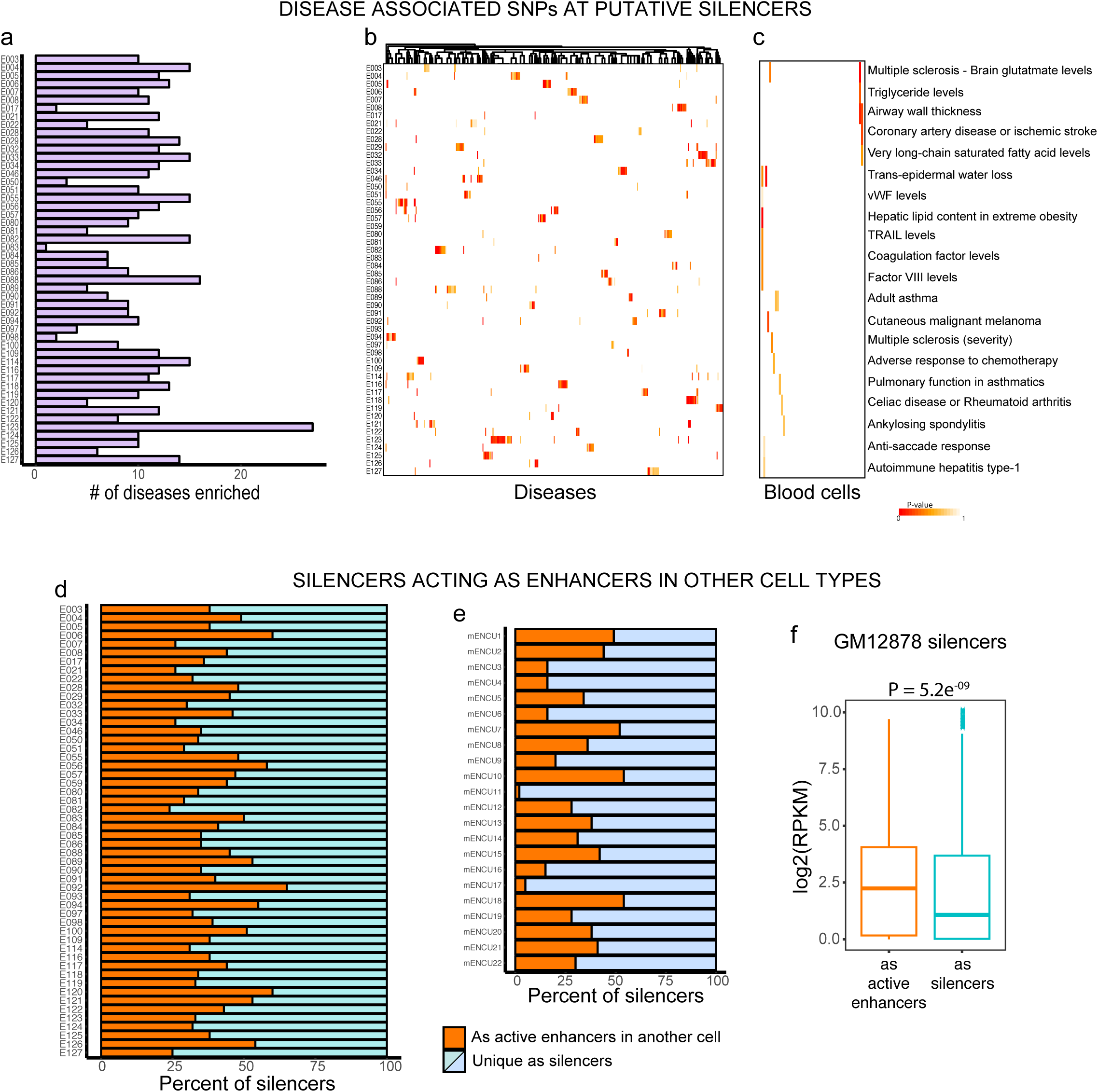
Enrichment of GWAS SNPs at putative silencer elements and putative silencers acting as enhancers in other cell types. **(a)** Bar plot presenting the count of GWAS SNPs belonging to various disease traits enriched (adj.P-value <0.01) at putative silencer elements of 52 cell types and tissues from Roadmap. (b) Heatmap of enrichment (-log10(P-value)) of 369 disease traits SNPs across 52 cell types. Columns represent disease traits and rows are cell types. (c) Heatmap of SNP enrichment (-log10(P-value)) for 20 autoimmune disease traits across 52 cell types. Bar plot showing the fraction of putative silencer elements identified in one cell type acting as enhancers in other cell types for human Roadmap data (d) and mouse ENCODE data (e). (f) An example of GM12878 silencers acting as enhancers in CD34+ cells. Box plot comparing the expression distribution of interacting genes.

### Silencers can act as enhancers in other cell types

Classic studies of promoter silencers showed that these elements reside next to activating sequences ^4^. The ability to repress expression can also extend to promoter-embedded enhancer elements. For example, an inducible enhancer within the promoter of the *IFNB* gene is held under negative control prior to induction by a proximal silencer element ^49^. We asked whether identified silencer elements in one cell type only act as silencers or can also enhance gene expression, i.e., act as active enhancers in other cell types. To verify this, we intersected the putative silencer elements from each cell type with active enhancers (overlapping DHS marked with H3K4me1 + H3K27ac) of the remaining cell types. Interestingly, we found that 25% to 65% of silencer elements from one cell type can act as enhancers in at least one other cell type (Fig. 6d). For the mouse silencer elements it varies from 1% to 55% (Fig. 6e). Next, we wanted to verify this feature of silencers acting as enhancers by looking at their interacting gene expression distribution. As a test case, we overlapped putative silencer elements in GM12878 cells with active enhancer elements of CD34+ cells. We then found enhancer interacting genes in CD34+ cells from p-CHiC data as was done for silencer interacting genes in GM12878 cells and compared the target gene expression levels in their respective cell types. The relative expression of silencer interacting genes in GM12878 cells is indeed significantly less compared to the expression of interacting genes in CD34+ cells when silencers are acting as enhancers (Fig. 6f). Promoter elements frequently harbor multiple, yet distinct, regulatory sequences, including the core promoter, enhancer elements and silencer elements. Our analysis of putative distal silencers elements indicates that a subset can likely function as or are part of the same *cis*-regulatory element as enhancers.

## DISCUSSION

The precise control of gene expression, either activation or repression, is essential for changes in cell fate and cellular response to external cues. That control is ultimately mediated by *cis*-regulatory elements in the genome and their cognate binding factors. Though silencers have shown to be analogues to enhancers in that they can be distal to the genes they regulate and often function in a position‐ or orientation-independent manner ^4^, silencers lack a unique chromatin signature to aid their genome-wide identification. Here, we developed a simple computational (SSA) approach to identify genome-wide putative silencer elements. We find that putative silencer elements are enriched with known repressor TFBS and motifs. P-CHiC interactions showed that inactive and lowly expressed genes are interacting with putative silencer elements. These results strongly support the annotation of silencer elements in human and mouse genomes. We acknowledge that this may still be an overestimate as we likely have not excluded all possible non-silencer *cis*-regulatory elements in our SSA predictions. These predictions also rely on the annotation of DHS in each genome, which are subject to some degree of both false positive and false negative calls.

In order to functionally validate a subset of SSA-predicted silencers, MPRAs were performed to test nearly 7500 putative elements. MPRA assays confirmed silencer activity for 51% of tested silencer elements. This validation rate is within the range or above what was shown for enhancer validation studies ^16,50^. Based on our MPRA data, we trained an SVM to refine our silencer predictions, which may serve as a more conservative estimate of elements in the respective genomes. However, it is susceptible to false negatives due to caveats of *in vitro* reporter assays. It is also feasible that weak silencers were not validated. The SVM classifier did improve the cellular specificity of putative silencer elements as has been shown for enhancers ^42–44^, which may aid future predictions when limited data are available.

The vast majority of disease-associated SNPs are known to occur outside of coding regions ^51^. Similar to enhancers, we found disease-associated SNPs are also enriched at putative silencers of relevant disease cell types or tissues ^17^. Mutations within silencer elements may provide one means for genes to escape repression in a disease-specific manner and have implications for other diseases such as various cancers.

Overall, we find that characteristics of the predicted putative silencers are expected of silencer elements and are in contrast to the activating activity of enhancers. The catalogue of putative silencer elements presented here across many cell types and tissues for human and mouse may serve as a resource that complements the ENCODE ^18^ and Roadmap consortia ^17^ catalogues for other *cis*-regulatory elements.

## ACKNOWLEDGEMENTS

R.D.H. is supported by funds from the NIH/NIAMS (R01AR065952), NIH/NIDDK (R01DK103667), and the United States-Israel Binational Science Foundation (2013027).

## AUTHOR CONTRIBUTIONS

N.D.J. and R.D.H. conceived and planned the study, and wrote the manuscript. N.D.J. performed all the analyses. A.J. performed the MPRAs. All authors reviewed the final manuscript.

## ONLINE METHODS

### Processed datasets used

We obtained uniformly processed consolidated epigenome data of ChIP-seq and DNase-seq from the Roadmap consortium for 52 human cell types. We used narrow peaks called by MACS2 program for DNase-seq and broad peaks called by MACS2 program for H3K4me1, H3K4me3 and H3K27ac histone modifications. We directly obtained the peaks files from NIH Epigenomics Roadmap project^17^ (http://egg2.wustl.edu/roadmap/web_portal/processed_data.html). Similarly, we obtained the data for mouse (mm10) cell types and another human 30 cell types from the ENCODE consortium (https://www.encodeproject.org/matrix/?type=Experiment) ^18^. We used the peak files generated by the uniform ENCODE Processing Pipeline. We merged the peaks files for each cell type if data are available from different sources. The CTCF TFBS are obtained from CTCFBSDB 2.0 (http://insulatordb.uthsc.edu) for both human and mouse ^52^. The gene TSS coordinates were obtained from GENCODE annotations for human (GRCh37.p13/hg19) and mouse (GRCm38.p5/mm10) ^53^.

### SSA approach

In the SSA approach we used DNase-seq peaks as open chromatin (DHS), H3K4me3 peaks as promoter elements, H3K4me1 peaks as enhancer elements and CTCF TFBS as insulator elements and active TSS are defined as 2000 bp upstream and 500 bp downstream of TSS of all GENCODE genes. We start with DHS peaks and removed DHS overlapping with any of H3K4me1 or H3K4me3 or active TSS or CTCF TFBS and the remaining non-overlapping DHS peaks are termed as putative silencer elements. We included CTCF TFBS from all cell types during subtractive analysis as TFBS are largely shared across cell types. We used bedtools suite for genomic subtractive analysis ^54^.

### Genomic annotation of putative silencers

The genomic annotations of predicted putative silencers are performed using HOMER suite (*annotatePeaks.pl*).

### Putative silencers overlap with repressor TFBS

The well-known repressor TFBS as REST, YY1, ZBTB33, SUZ12 and EZH2 for cell types GM12878, H1, K562 and HEPG2 are directly downloaded from ENCODE project as processed peaks file. We computed the enrichment (number of overlaps) of repressor TFBS at putative silencer elements using bedtools suite.

### Motif enrichment analysis

Motif analysis to find enriched TF motifs at putative silencer elements were performed with HOMER suite (*findMotifsGenome.pl*‐size given). Enriched motifs are defined at adjusted P-value < 0.001. Heatmap visualizations are created using R package ‘pheatmap’.

### Computing average methylation values

The methylation value for each cytosine from WGBS data for 17 cell types were obtained from NIH Epigenomics Roadmap project as bigwig files. The bigwig files were converted to bedgraph files via UCSC file conversion tool, and we computed the average methylation levels across putative silencer elements and active enhancers.

### chromHMM annotation datasets

The chromHMM annotation categories for all 52 human cell types are obtained from NIH Epigenomics Roadmap project. We used the core 15-state model for chromatin state learning for predicted putative silencer elements and for TSS of genes interacting with silencer elements.

### Computing average phastCon conservation scores

We computed average phastCons scores for putative silencer elements and active enhancers using R/Bioconductor package, ‘phastCons100way.ucsc.hg19’ with ‘scores’ function.

### Promoter capture HiC datasets used

We obtained promoter capture HiC data for GM12878 and CD34+ cells in human from ^30^ and mouse embryonic cells (mESCs) and mouse fetal liver cells (FLCs) from ^31^. The processed significant promoter-other end fragments interactions were directly obtained from the authors supplementary data. The corresponding promoter gene expression data for GM12878 and CD34+ cells are obtained from NIH Epigenomics Roadmap project and for mESC and FLC are obtained from the ENCODE project.The putative silencer elements and active enhancer elements interacting genes were obtained by overlapping respective *cis*-regulatory elements with other end fragments. We consider a minimum of 1 bp overlap. We defined active enhancers as DHS overlapping with both H3K4me1 and H3K27ac (DHS + H3K4me1 + H3K27ac).

### Cell Culture

All experiments are performed in K562 cells. Cells obtained from the ATCC(CCL-243) are grown at 37°C and 5% CO2 in RPMI-1640 (Gibco) medium with 10% fetal bovine serum (Gibco) and 1% penicillin-streptomycin.

### Massively Parallel Reporter Assays (MPRAs)

To multiplex silencer validation assay, we leveraged synthetic oligonucleotide array synthesis and adopted Self-transcribing active regulatory region sequencing (STARR-seq) method as described in (Arnold et al, Science.2013). An oligonucleotide library was synthesized containing 200nt of genomic regions with 15 nt of flanking sequence matching the Illumina primer sequence (Agilent, Inc). The obtained library was PCR amplified to add Illumina primer sequence and 15 bp of sequence matching the STARR-seq backbone for In-Fusion cloning (Clontech). Transfection were performed in K562 cells ATCC(CCL-243) using Neon transfection kits and device (Invitrogen) as per manufacturer’s recommendations. Total RNA was prepared using Qiagen RNeasy Plus Mini kit and Poly A-RNA was isolated from total RNA using μMACS mRNA Isolation Kit (Miltenyi Biotech) and and plasmid DNA were extracted by Qiagen Plasmid Plus Midi kit. All libraries were sequenced in NextSeq 500 (Illumina) performing 1 × 75 cycles.

### MPRA data normalization and analysis

Sequencing raw reads from RNA and plasmid libraries are checked for adapter sequences and low quality reads (q score < 20) using FASTQC (https://www.bioinformatics.babraham.ac.uk/projects/fastqc) and trimmed using TrimGalore package (https://www.bioinformatics.babraham.ac.uk/projects/trim_galore/). Trimmed reads were mapped to human genome (hg19) using the Bowtie2 aligner ^55^. The mapped reads were quantified against all tested sequences using ‘featureCounts’ function from Rsubread package ^56^. The biological replicates were checked for similarity and pooled to increase the statistical power. The RNA and plasmid read counts were normalized by respective library size and the normalized regulatory element activity is defined as ratio of RNA to plasmid counts averaged over biological replicates for each tested sequence. Fisher’s exact test was used to find significant differences in tested silencer activity over control random regions activity. The computed P-values are adjusted with Benjamini-Hochberg method to control for false discovery rate. Then, the potential silencers were identified as sequences with activity less than median activity of control random regions and adjusted P-value < 0.1.

### SVM model

We used the R package ‘gkmSVM’ for SVM model training and putative silencer predictions in other cell types ^45^. We chose the top 2200 putative silencer sequences with lowest activity as positive set and bottom 2200 silencers with highest activity as negative set for SVM model training with default settings from package. We used 80% of data for training and the remaining 20% data for testing the model. We evaluated the performance of the model on test data by generating the receiver operating characteristic (ROC) curve by plotting true positive rate versus false positive rate and the precision recall curves (PRC).

### Disease SNPs analysis

Disease-associated SNPs were obtained from NHGRI GWAS catalog (https://www.ebi.ac.uk/gwas/docs/file-downloads) and SNP coordinates are converted from GRCh38 to hg19 using UCSC liftOver tool. We included both lead SNPs and SNPs in LD block (r^2^ >= 0.8) for overlap with silencer elements. The SNPs in LD (proxy SNPs) were obtained from SNAP webserver (http://archive.broadinstitute.org/mpg/snap/ldsearch.php) ^57^. The enrichment of disease SNPs to putative silencer elements for each cell type are obtained by comparing the relative enrichment of particular disease SNPs against the rest of disease SNPs using a hypergeometric distribution.

### Statistical tests and visualizations

All the statistical tests are performed in the R environment (https://www.r-project.org). Graphs are prepared using ‘ggplot2’ and ‘gplots’ R packages.

## Supplemental information

**Supplementary Table 1: Overview of cell types and count of putative silencer elements per cell type**

**Supplementary Table 2: MPRA tested silencer elements along with control elements with their activity levels.**

**Supplementary Table 3: Overview of SVM classifier identified putative silencer elements**

**Supplementary Figure 1.**
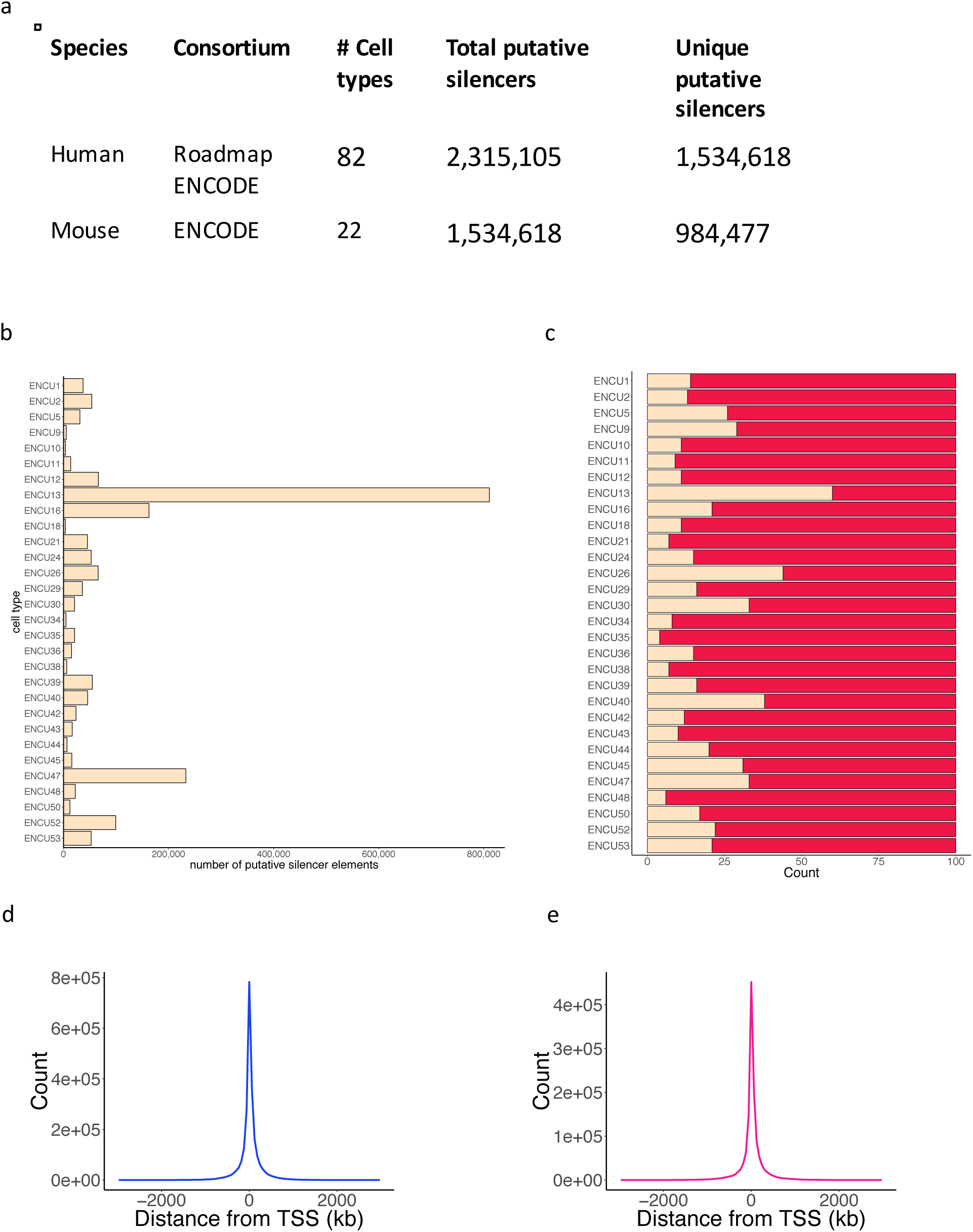
Overview of identified putative silencer elements. **(a)** Count of total and cell type specific putative silencer elements identified across human and mouse genomes. (b) Bar plots presenting the count of predicted silencer elements across different cell types and tissues in human from ENCODE. (c) Distribution of cellLtype specific silencer elements in human from ENCODE cell types. The location of putative silencers relative to genes TSS in human (d) and mouse (e)

**Supplementary Figure 2.**
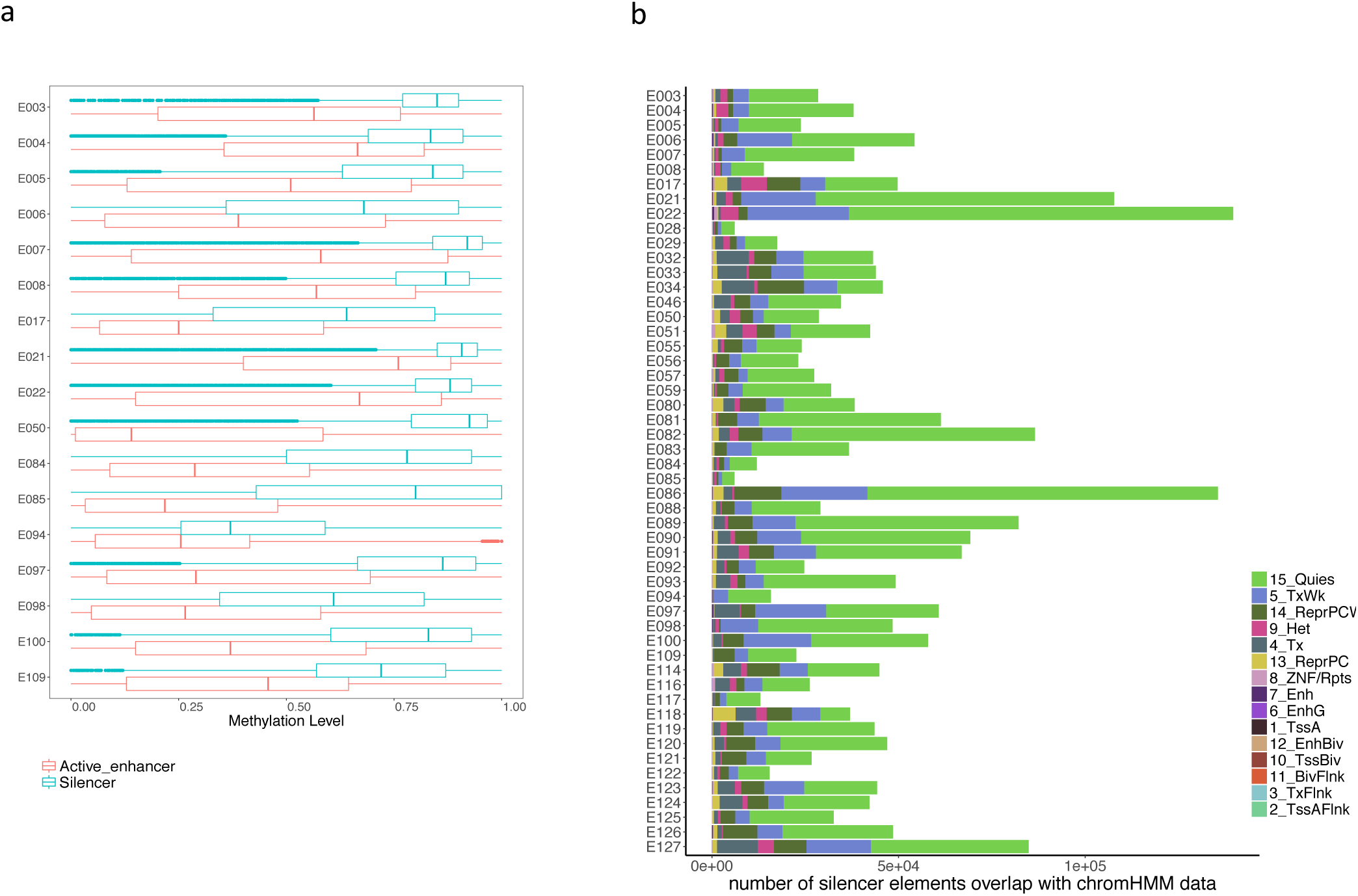
Characteristics of putative silencer elements. **(a)** Box plot showing the distribution of methylation levels at putative silencer elements and active enhancers across cell types. (b) Bar plot showing the distribution of chromHMM annotations of putative silencer elements across 52 cell types in human.

**Supplementary Figure 3.**
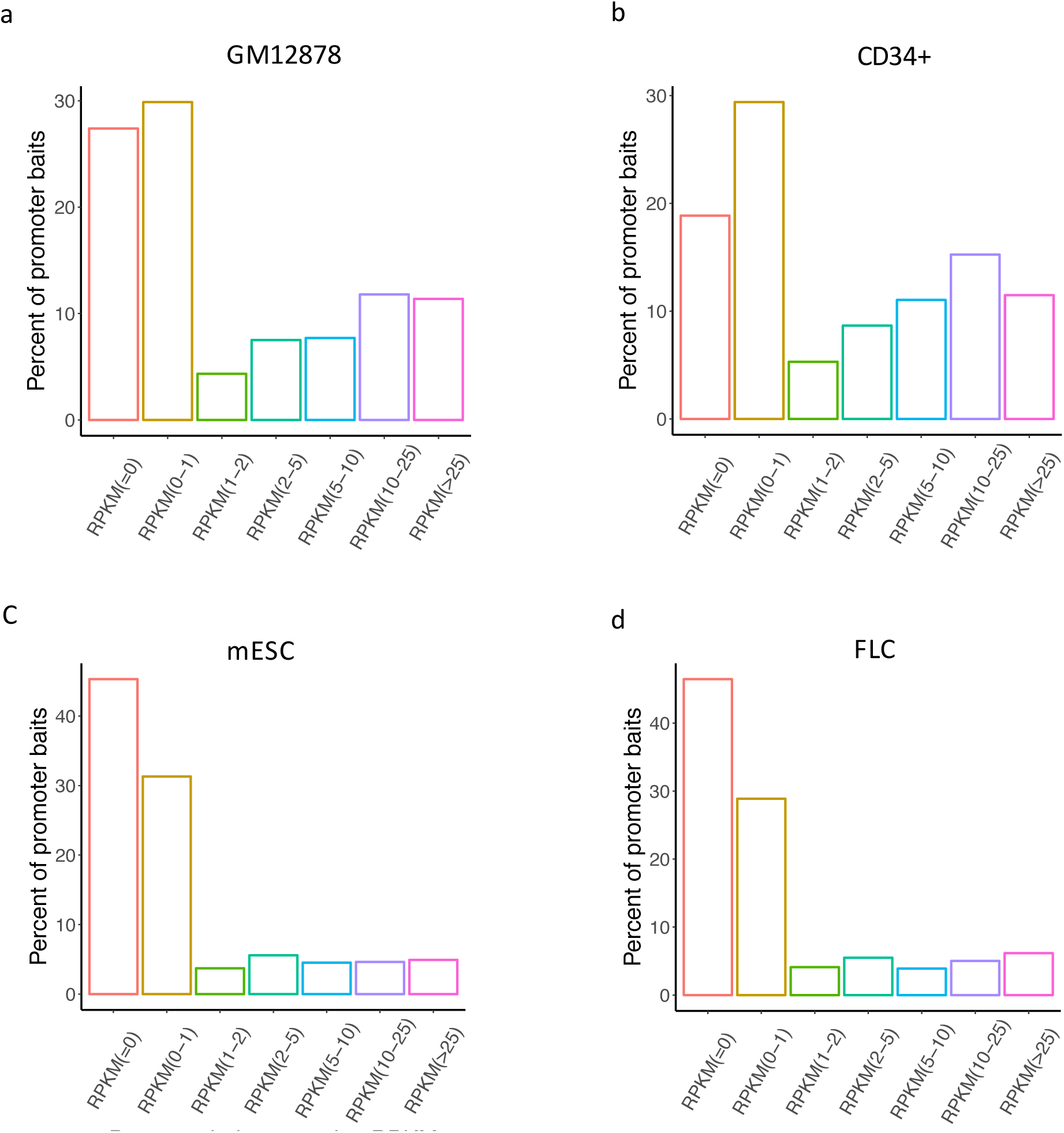
3D Genome interactions of putative silencer elements. Bar plot presenting the percent of total genes expressed at different expression levels (RPKM) interacting with putative silencer elements in GM12878 cells (a), CD34+ cells (b), mESCs (c), mouse fetal liver cells (d).

**Supplementary Figure 4.**
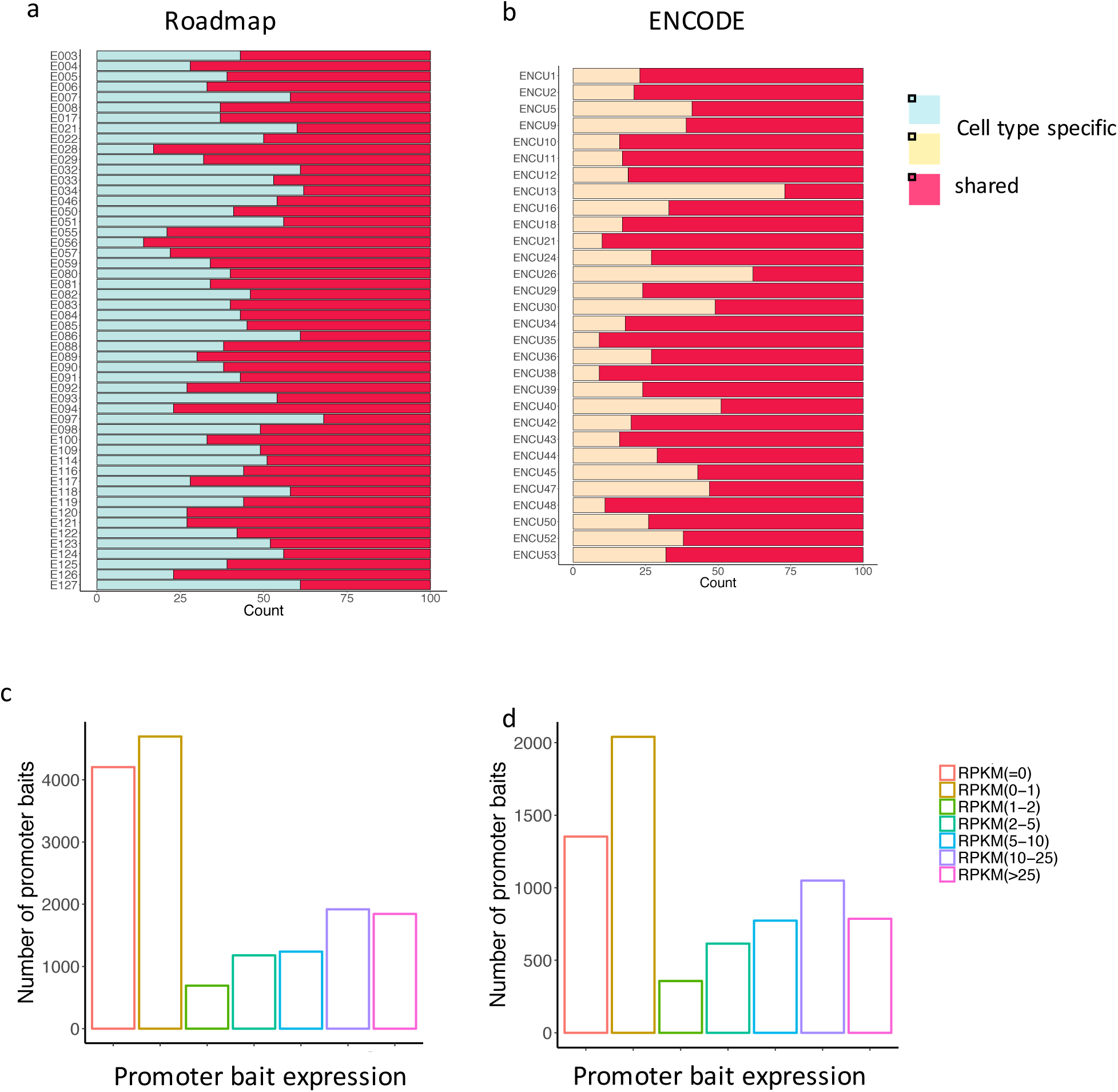
Overview of SVM model and silencer elements predictions. Distribution of SVM predicted cellLtype specific silencer elements in human from Roadmap (a) and ENCODE (b). Bar plot presenting the count of genes expressed at different expression levels (RPKM) interacting with SVM predicted silencer elements in GM12878 cells (c), CD34+ cells (d).

## SUPPLEMENTAL EXPERIMENTAL PROCEDURES

### Designand Synthesis of a STARR-seqLibrary

To multiplex silencer validation assay, we leveraged synthetic oligonucleotide array synthesis and adopted selfLtranscribing active regulatory region sequencing (STARRLseq) method. An oligonucleotide library was synthesized containing the 230 nucleotide genomic regions with 15 bp of sequence matching the Illumina primer sequence was performed by Agilent Inc. The obtained library was then PCR amplified to add Illumina primer sequence and 15 bp of sequence matching the STARRLseq backbone. The library was amplified with KAPA Hi‐fi 2X. (Kapa biosystem) with following thermal condition (98 °C for 2 min, amplification with ten cycles of 98 °C for 20 s, 65 °C for 15 s, and 72 °C for 30 s; final extension at 72 °C for 2 min). The resulting product was purified using Ampure XP beads at 1.8X beads: reaction ratio. The STARRLseq screening vector digested for six hour with SalILHF and AgeILHF and linearized backbone run on the gel was purified with the Gel purification kit (Qiagen). 200 ng of backbone and 50 ng pooled insert was cloned in four 10 μl InfusionLHD reactions incubating at 50°C for 15 min (Clontech). Resulting products were then combined purified using AmpureLXP beads with a 1X volume of beads and eluted in 8 μl of purified water and electroporated into NEB^®^ 10Lbeta electro competent cells at a ratio of 2 μl of reaction to 20 μl of competent cells for a total of four electroporations using BioLRad GenePulserR II electroporator, with following electroporation conditions: 2.0 kV, 200 Ω, 25 μF. Transformations were recovered for one hr in SOC medium while shaking (220 rpm, 37 °C) and then grown for 16 hr in 500 mL of Luria Broth while shaking (220 rpm, 37 °C). The STARRLseq input libraries were then purified using the Qiagen Plasmid Maxiprep kit. Cells were electroporated with a Neon transfection kit and device (Invitrogen) with following transfection parameters, Pulse voltage: 1450 Pulse width: 10 Pulse number: 3.

### Output Library Construction

Cell pellets were rinsed once with PBS and then lysed in 2 ml of RLT buffer (Qiagen) with 2L mercaptoethanol (Sigma).Total RNA was prepared using the QIAGEN RNeasy kit. PolyLA RNA isolated from 50 μg of total RNA by μMACS mRNA isolation kit (Miltenyi Biotech). RNA was then treated with turboDNase (4 U) for 30 min at 37°C (Invitrogen). DNase treated polyLA RNA was purified using the RNeasy kit. Plasmid specific cDNA was synthesized using Superscript III (Life Technologies) incubated for 1.5 hr at 55°C and inactivated at 80°C for 15 min. Following synthesis, cDNA was treated with RNaseA (Sigma) at 37°C for 30 min. cDNA was purified using Ampure beads in 1.5:1 beads:cDNA ratio and then amplified and indexed for sequencing using a twoLstage PCR as described previously (Arnold et al. Science Reference). The cDNA sample from each replicate was used as an input into first round geneLspecific PCR reaction using primers “AJ10Lsequence” and “AJ12Lsequence”, and KAPA hiLfidelity polymerase (KAPA biosystem). PCR conditions were (98 °C for 2 min, amplification with 15 cycles of 98 °C for 20 s, 65 °C for 20 s, and 72 °C for 60 s; final extension at 72 °C for 2 min).

Samples were then purified using Zymo PCR purification kit and eluted in 15 μl nuclease free water. The resulting products were used as template for the second round of PCR, which used a standard Illumina TruSeq indexing primer on the p5 end of the library and custom indexing primers (sequences to be added in Suppli table) to barcode the samples for multiplexing prior to sequencing (98 °C for 2 min, amplification with 8 cycles of 98 °C for 15 s, 65 °C for 30 s, and 72 °C for 30 s; final extension at 72 °C for 2 min. Final sequencing libraries were purified with Ampure XP beads (Beckman Coulter) at a 1.8x SPRI: PCR reaction ratio. All libraries were sequenced in NextSeq 500 (Illumina) performing 1 × 75 cycles.

